# Simulating Longitudinal Single-cell RNA Sequencing Data with RESCUE

**DOI:** 10.1101/2025.01.06.631629

**Authors:** Elizabeth A. Wynn, Kara J. Mould, Brian E. Vestal, Camille M. Moore

**Affiliations:** Center for Genes, Environment and Health, National Jewish Health, Denver, CO, USA; Department of Medicine, National Jewish Health, Denver, CO, USA; Department of Pulmonary and Critical Care Medicine, University of Colorado Anschutz Medical Campus, Aurora, CO, USA; Department of Biostatistics and Informatics, University of Colorado Anschutz Medical Campus, Aurora, CO, USA

## Abstract

As single-cell RNA-sequencing (scRNA-seq) becomes more widely used in transcriptomic research, complex experimental designs, such as longitudinal studies, become increasingly feasible. Longitudinal scRNA-seq enables the study of transcriptomic changes over time within specific cell types, yet guidance on analytical approaches and resources for study planning, such as power analysis, remains limited. Data simulation is a valuable tool for evaluating analysis method performance and informing study design decisions, including sample size selection. Currently, most scRNA-seq simulation methods simulate cells for a single sample, thus ignoring the between-sample and between-subject variability inherent to longitudinal scRNA-seq data. Here, we introduce RESCUE (REpeated measures Single Cell RNA-seqUEncing data simulation), a novel method that simulates longitudinal scRNA-seq data using a gamma-Poisson framework and incorporates additional variability between samples and subjects. We demonstrate our method’s ability to reproduce important data properties and demonstrate its application in study planning. RESCUE is implemented as an R package and is available at https://github.com/ewynn610/RESCUE.

## 1 Introduction

RNA-sequencing (RNA-seq) has revolutionized our understanding of how disease and other biological conditions impact gene expression by simultaneously measuring the expression levels of thousands of genes across the transcriptome [Ozsolak and Milos, 2011, Stark et al., 2019]. Using traditional “bulk” RNA-seq, messenger RNA from all cells in a sample, which may include several distinct cell types, is sequenced together. The resulting expression represents an average across all cells in a sample, and differences in expression between conditions may be due to differences in cell type composition. Single-cell RNA-sequencing (scRNA-seq) technologies have addressed this limitation by enabling the sequencing of RNA from individual cells, eliminating the confounding effects of cell type composition. This has allowed for several novel applications, including the investigation of differential gene expression between biological conditions or disease states for a specific cell type or subpopulation of cells.

With the decreasing cost of scRNA-seq, longitudinal and paired study designs have become feasible, allowing researchers to sequence cells from multiple time points or conditions within the same individual. After processing and integrating the data, cells are typically categorized into specific cell types or subpopulations of cells (Fig. 1a). The repeated measures approach then enables the study of temporal changes in the transcriptome within particular cell types or subpopulations. Although such studies are currently being implemented [Maynard et al., 2020, Ostasov et al., 2020, Kazer et al., 2020, Kim et al., 2018], there has yet to be a comprehensive assessment of the best methods to analyze this type of data. Additionally, due to the novelty of longitudinal scRNA-seq studies, there are few resources to guide study design.

**Figure 1:**
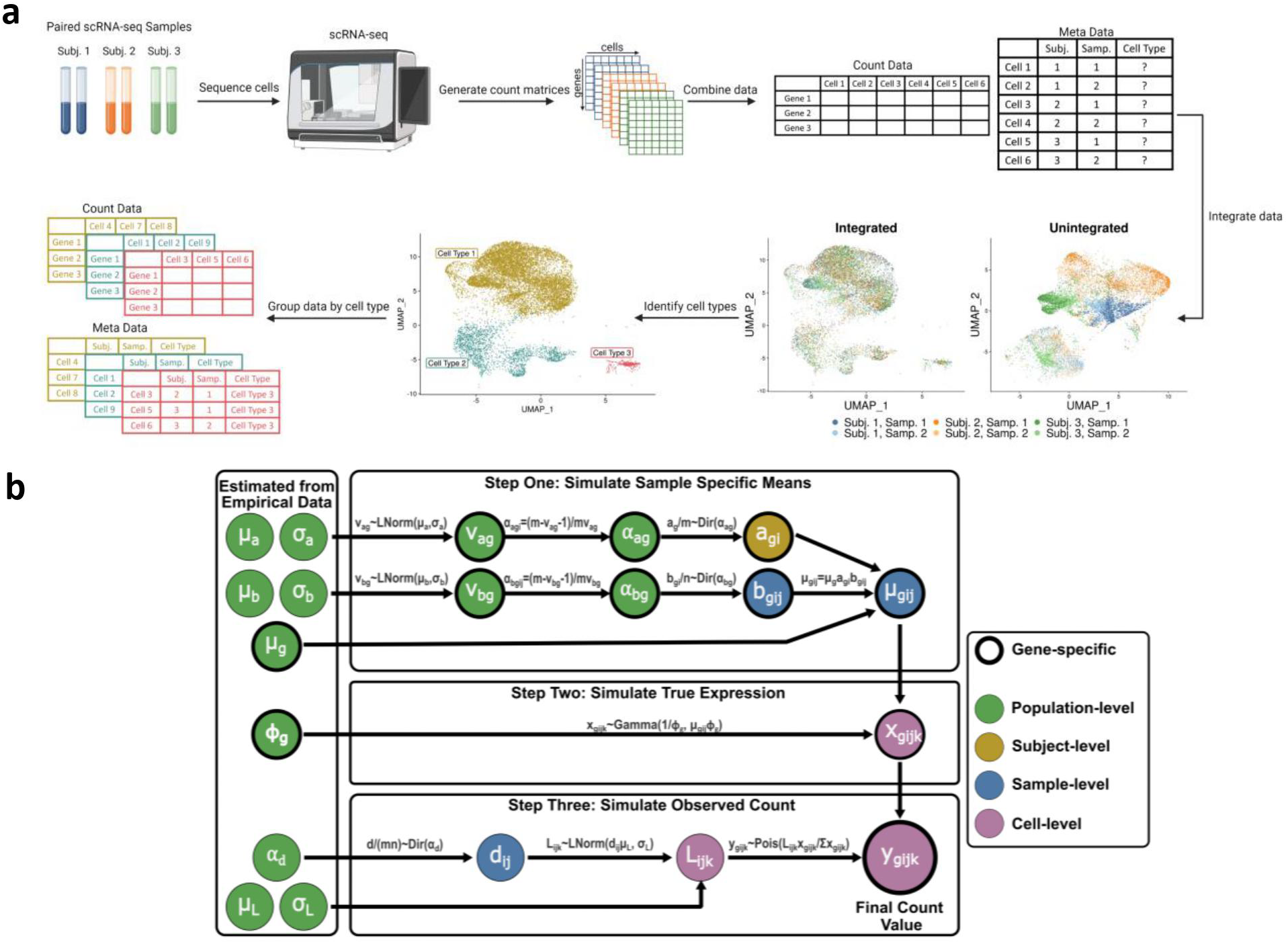
a) Summary of a typical data preparation workflow for longitudinal scRNA-seq data. Longitudinal samples are sequenced, the resulting count matrices are combined and the data are integrated, cell types are identified through clustering, and count data is separated by cell type. b) Overview of the simulation framework. Parameters and estimates are shown in circles, with the colors indicating whether they are population, subject, sample, or cell-level values. Circles with bold outlines represent parameters estimated at a gene level. The left most values are estimated from empirical data.

The ability to simulate scRNA-seq data for complex study designs could aid in selecting appropriate analysis methods and inform study design decisions. Simulating data allows researchers to control key data characteristics, enabling them to evaluate and compare the accuracy of statistical methods across various scenarios. Data simulation can also be useful for planning complex studies where closed-form formulas for power and sample size calculations may not be available. For example, researchers can simulate datasets with different sample sizes and assess power to inform sample size selection.

ScRNA-seq data are typically large and complex, encompassing information from thousands of genes across thousands of cells, making data simulation challenging. There are many properties that characterize scRNA-seq datasets including the number of cells, cellular library size, average expression levels, and proportion of zero-count observations. One particularly complex aspect of longitudinal scRNA-seq data is its hierarchical structure, where cells are nested within each sample, and multiple samples exist for each subject. This structure introduces both between-sample and between-subject variability, which must be accounted for in analysis and study planning. This variability arises from both biological and technical factors. Biological differences between samples and subjects can cause gene expression in cells from the same sample or from the same subject at different timepoints to be more similar than in cells from different samples or subjects. Additionally, cells from a single sample are usually processed and sequenced together, which can introduce a technical batch effect.

Several simulation methods have been developed for cross-sectional scRNA-seq data. However, most of these methods only simulate data from a single sample, which is generally inadequate for experiments that include multiple biological replicates. Some simulation packages offer options to simulate batch effects [Baruzzo et al., 2020, Zappia et al., 2017, Assefa et al., 2020, Zhang et al., 2019]. By treating cells from individual samples as batches, these methods can be adapted to simulate multi-sample data. However, the performance of the batch effect modifications to single sample simulation methods has not been fully evaluated in the literature, and the modifications are not always thoroughly described in the associated R package documentation. The only well-developed method for simulating multi-sample scRNA-seq data is the `splatPop` method [Azodi et al., 2021]. This method uses a gamma-Poisson framework to simulate counts, incorporating a sample-specific mean for each gene to ensure that expression levels of cells from the same sample are correlated. Currently, there are no methods available that can simulate the hierarchical structure of longitudinal scRNA-seq data, which includes both between-sample and between-subject variability.

In this paper, we propose the REpeated measures Single Cell RNA-seqUEncing data simulation (RESCUE) method for simulating longitudinal scRNA-seq data. This method employs a gamma-Poisson framework to simulate counts, incorporating additional variability between cells from different samples and subjects to replicate the hierarchical structure of longitudinal data. We demonstrate that our method can reproduce several key data properties observed in empirical longitudinal scRNA-seq data from various cell types. Additionally, we show how RESCUE can be utilized for sample size selection and power analysis in study planning. RESCUE is implemented as an R package, available for installation at https://github.com/ewynn610/RESCUE.

## 2 Simulation Framework

Our simulation method generates a longitudinal scRNA-seq dataset for a single cell type or sub-population of cells, providing gene expression values for *m* subjects, each with *n* samples, and an average of *c* cells per sample, where the user can specify the values for *m, n*, and *c*. For multiple cell types of interest, the simulation process can be repeated for each cell type. Empirical data for each cell type is required and can be obtained from pilot data or from a publicly available database such as the Gene Expression Omnibus [Edgar et al., 2002]. Multi-sample or longitudinal data is not required.

The simulation framework consists of three steps (Fig. 1b). First, we estimate sample-specific mean expression values, with correlation between values for samples from the same subject. Second, using these mean values, we draw “true” expression values for each cell. Finally, we simulate final count values for each cell, incorporating additional variability to represent technical variation. Similar to Baruzzo et al. [2020], the *true* gene expression value for gene *g*, subject *i*, sample *j* and cell *k*, denoted as *x*_*gijk*_, is unknown in an empirical dataset, while the *observed* expression value, denoted as *y*_*gijk*_, can be considered the observed count value from an empirical dataset.

### 2.1 Step One: Simulating Sample-specific Means

#### 2.1.1 Drawing Simulated Values

Let *µ*_*g*_ be the global mean, representing the average expression of gene *g* across all samples. Due to technical and biological factors, the mean expression of gene *g* for subject *i* and sample *j, µ*_*gij*_, will differ from *µ*_*g*_. To account for between-subject and between-sample variation in mean expression levels, we calculate *µ*_*gij*_ = *µ*_*g*_*a*_*gi*_*b*_*gij*_, where *a*_*gi*_ and *b*_*gij*_ are multiplicative factors that allow for subject- and sample-specific deviations from the global mean, respectively.

We draw *a*_*gi*_ and *b*_*gij*_ from symmetric Dirichlet distributions. Let **a**_**g**_ = {*a*_*g*1_, …*a*_*gm*_} and **b**_**gi**_ = {*b*_*gi*1_, …, *b*_*gin*_}. Then

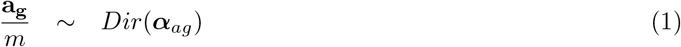

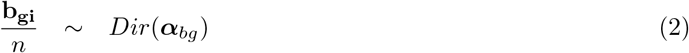

where ***α***_*ag*_ and ***α***_*bg*_ are gene-specific concentration parameters.

To obtain values for ***α***_*ag*_ and ***α***_*bg*_, we draw *v*_*ag*_ and *v*_*bg*_ from log-normal distributions:

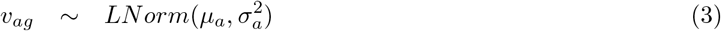

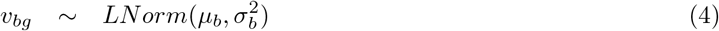

where *v*_*ag*_ and *v*_*bg*_ are the variances of *a*_*g*_ and *b*_*gi*_, respectively. The hyper-parameters *µ*_*a*_ and *µ*_*b*_ represent the log-scale mean values for *v*_*ag*_ and *v*_*bg*_ while 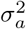 and 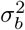 represent the log-scale variances. We use the relationship between the variance and concentration parameter to calculate ***α***_*ag*_ and ***α***_*bg*_:

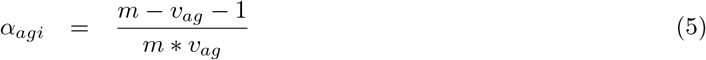

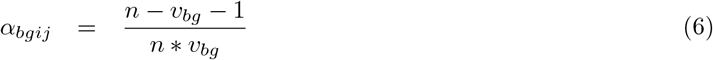

#### 2.1.2 Estimating Hyper-Parameters

We estimate hyper-parameters from empirical data by first normalizing the data using the scran normalization method [Lun et al., 2016]. For each sample, we then estimate the gene-specific mean value using 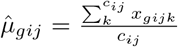, where *x*_*gijk*_ represents cell-level, gene-specific normalized expression values and *c*_*ij*_ is the number of cells from the sample. The overall gene-specific mean, *µ*_*g*_, is then estimated as 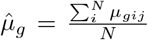, where *N* is the total number of samples in the empirical dataset.

The estimation of parameters *µ*_*b*_, *σ*_*b*_, *µ*_*a*_, and *σ*_*a*_ requires multi-sample data (for *µ*_*b*_ and *σ*_*b*_) or longitudinal data (for *µ*_*a*_ and *σ*_*a*_). If such data is unavailable, we provide plausible values estimated from the datasets described in Section 3.1.1 (Supplementary Table 1). When estimating these parameters from empirical data that include samples from different conditions or timepoints, the parameters should be estimated using a set of genes that remain unchanged across conditions/timepoints or after regressing out condition/timepoint effects (Supplementary Material section 1).

To estimate *µ*_*b*_, *σ*_*b*_, *µ*_*a*_, and *σ*_*a*_ from empirical data, we calculate the batch effects *a*_*gi*_ and *b*_*gij*_ from the empirical data using the following equations:

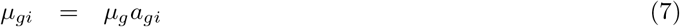

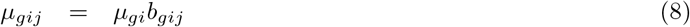

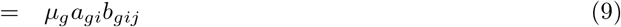

Once we have generated sample and subject level batch factors for each gene, we take the variance such that:

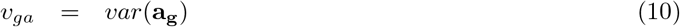

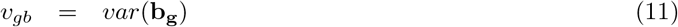

Because we calculate *a*_*gi*_ and *b*_*gij*_ using sample level means, we would expect some level of variation due to sampling error even if there were no true variation in the sample or subject level means. To account for this, we estimate the gene specific error variance and subtract it from the total variance to get the variance due to between sample/subject variability,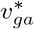 and 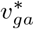 (Supplementary materials section 1).

Finally, to obtain the parameters, *µ*_*b*_, 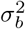, *µ*_*a*_, and 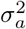, we take the mean and variance across genes For 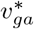 and 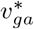. Then, using the relationship between the mean/variance and the log mean and variance parameters of the log-normal distribution, we calculate *µ*_*b*_, *σ*_*b*_, *µ*_*a*_, and *σ*_*a*_. We found that for genes with a high percentage of 0’s, the estimates for 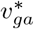 and 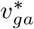 are unstable, so we only use genes with *<* 60% 0 counts to estimate the parameters.

### 2.2 Step Two: Simulating True Expression Values for Each Cell

#### 2.2.1 Drawing Simulated Values

Using the sample-specific means estimated in step one, we draw “true” expression values for each cell from a gamma distribution:

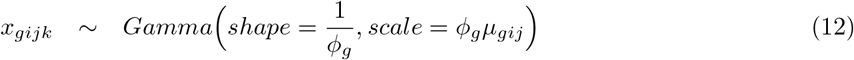

where *ϕ*_*g*_ is a gene-specific dispersion parameter.

#### 2.2.2 Estimating Hyper-Parameters

Values of *ϕ*_*g*_ are calculated from the empirical data using the `estimateDisp` function in the `edgeR` package [Robinson and Oshlack, 2010]. If multi-sample data is provided, the sample with the most cells is used.

### 2.3 Step Three: Simulating Observed Counts for Each Cell

#### 2.3.1 Drawing Simulated Values

Before simulating the observed counts, we draw an expected library size for each cell from a log-normal distribution:

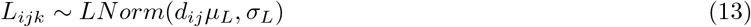

where *µ*_*L*_ and *σ*_*L*_ are the mean and standard deviation of the log-library sizes and *d*_*ij*_ is used to simulate sample-specific differences in the average library size due to technical batch effects. Let **d** = {*d*_11_, …, *d*_*mn*_}.

Then we use a symmetric Dirichlet distribution to generate **d**:

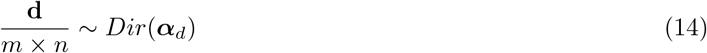

where *m* × *n* represents the total number of samples being simulated and ***α***_*d*_ is a vector of identical concentration parameters of length *m* × *n*.

Using the expected library size values, we scale the simulated true expression values using 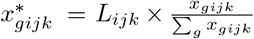 where 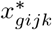 is the library-adjusted value for *x*_*gijk*_. Then, using these values, we simulate our final observed count values for each cell using a Poisson distribution:

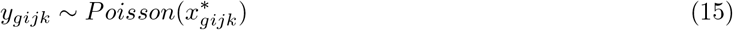

#### 2.3.2 Estimating Hyper-Parameters

The *µ*_*L*_ and *σ*_*L*_ parameters are calculated by taking the sample-specific mean and standard deviation of the log library sizes for each sample from the empirical data and then averaging the values across all of the samples.

The concentration parameter ***α***_*d*_ is estimated by calculating the deviation between the global average log library size and the sample-specific averages, calculating the variance between these deviations, and then generating a parameter estimate using the relationship between the variance and the symmetric Dirichlet concentration parameters previously outlined.

### 2.4 Simulating Differential Expression

A slight adjustment to step two of our framework can be made to simulate differential expression between timepoints. Suppose we have two different timepoints (or biological conditions/disease states), A and B. For the cells in timepoint A, we simulate the true expression value as previously outlined. For timepoint B, we will simulate the true expression values of *N* genes to be differentially expressed from timepoint A, with an expected fold change of *z*_*g*_, where *z*_*g*_ > 0. Thus,

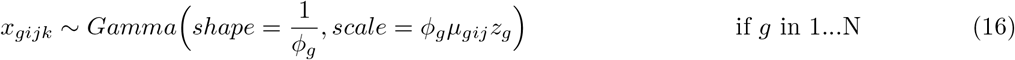

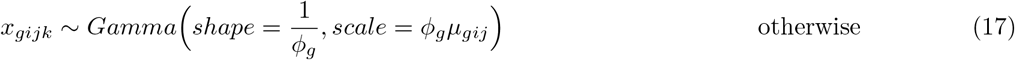

The fold change values are provided by the user as a vector with one value per gene. Each value represents the fold change value for a given gene, where a value of one indicates no differential expression, values below one represent downregulation at timepoint B relative to timepoint A, and values above one represent upregulation at timepoint B relative to timepoint A.

## 3 Methods

### 3.1 Simulation Assessment

#### 3.1.1 Assessment Datasets

We assessed our simulation framework using two empirical scRNA-seq datasets. First, we used five samples from a dataset containing bronchoalveolar lavage (BAL) samples from healthy adult subjects without any treatment or intervention (GEO accession: GSE151928) [Mould et al., 2020]. To calculate parameters relating to between-subject variability, we used data provided by the study authors containing samples for each subject 4-5 days after administration of lipopolysaccharides (LPS), which was used to induce acute lung inflammation. Before simulation, the standard Seurat workflow was used to integrate cells from different samples, cluster cells using a shared nearest neighbor algorithm, and identify markers specific to each cluster for cell type assignment [Stuart et al., 2019] (Fig. 1A).

The second dataset contained blood samples from six children with mild/asymptomatic COVID-19 [Khoo et al., 2023]. We used the processed Seurat dataset available on the GEO database (accession: GSE196456) with the cell types annotated using the Azimuth algorithm in Seurat [Hao et al., 2021]. For each dataset, we simulated data for cell types with at least 25 cells in each sample, resulting in four simulated cell types for the Khoo et al., 2023 dataset and three for the Mould et al., 2020 dataset. For each cell type, we removed genes with no counts across all cells and simulated paired data for the same number of subjects as in the empirical dataset. The number of cells per sample was randomly drawn from a discrete uniform distribution using the minimum and maximum cells per sample for each timepoint from the empirical data as parameters. All data were simulated without differential expression. For brevity, we focus our results on two cell types, the resident airspace macrophage (RAM) cells from the Mould et al., 2020 dataset and the B cells from the Khoo et al., 2023 dataset. Results from other cell types are available in Supplementary material Figs. 1-5.

#### 3.1.2 Assessment Metrics

We compared the distributions of key metrics from the simulated and empirical data, including the mean and variance of log counts per million normalized gene expression levels, proportion of zero-counts across genes and cells, and the cellular library size. To assess our method’s ability to recreate the hierarchical structure of the empirical data, we compared sample- and subject-level clustering in t-distributed stochastic neighbor embedding (tSNE) plots from unintegrated simulated and empirical data. We also compared the distribution of intra-class correlation (ICC) values at the subject and sample levels. To calculate ICC, we first transformed the data using a variance stabilizing transformation from the `sctransform` R package [Hafemeister and Satija, 2019]. Then, using the `lmerSeq` package [Vestal et al., 2022], we fit linear mixed models for the 800 genes with the highest counts per million (CPM) values. These models included a fixed effect intercept and random intercepts for sample and subject. From these models, we calculated the ICC values for sample and subject. tSNE and ICC values were calculated using the set of invariant genes found during simulation (see Supplementary material section 1).

### 3.2 Power Analysis

To demonstrate the utility of our method, we performed a hypothetical power analysis for a scRNA-seq study investigating differential expression between timepoints in RAM cells. Specifically, we explored whether greater gains in power are obtained by recruiting more subjects, gathering data from more timepoints per subject, or sequencing more cells.

We used the RAM empirical data to simulate 10 datasets for each of four scenarios. The baseline scenario included five total subjects, two timepoints per subject, and an average of 200 RAM cells per sample. In the remaining scenarios, either the number of subjects, timepoints, or cells was doubled. To generate the data for each scenario, we simulated data with 10 subjects, four timepoints, and 400 cells, then down-sampled the subjects, samples, or cells according to the specifications of each scenario. We randomly drew the number of cells per sample from a uniform distribution with the minimum and maximum values set to *±*100 the desired average value. Twenty percent of genes were simulated to have a log_2_ fold change of *±*0.35 between the first and final timepoint. In the simulation with four timepoints, we simulated a linear change in expression for differentially expressed genes.

We used the `MAST` package to assess differential expression in each simulated dataset [Finak et al., 2015]. This package employs a two-step hurdle model on normalized data to test for differential expression, with a logistic regression model to determine whether the gene is expressed, and a linear model to assess gene expression levels (given the gene is expressed). This model is one of the only scRNA-seq specific methods that allows for multiple random effects, making it suitable for longitudinal data. Before running MAST, we filtered out genes with more than 90% zero-counts from each simulated dataset and applied the *log*_2_ + 1 transcripts per million transformation suggested in the MAST documentation. We included random intercepts for sample and subject and fixed effects for time and the number of genes detected in a cell, as recommended by Finak et al. [Finak et al., 2015]. Likelihood ratio tests using the combined hurdle method were used to test for differential expression across time. P-values were adjusted using the Benjamini-Hochberg method [Benjamini and Hochberg, 1995], and we assessed differential expression at three significance thresholds, 0.01, 0.05, and 0.1. Power was calculated for each dataset at each threshold.

## 4 Results

### 4.1 Simulation Assessment

Our simulated data reproduced important properties from empirical data. The average and variance of normalized gene expression closely matched between empirical and simulated data, showing correlation values > 0.99 for both RAM and B cells (Fig. 2a-b). Additionally, the simulated data successfully captured the gene-level relationship between average expression and variance (Fig. 2c). RESCUE also effectively replicated data sparsity, as evidenced by the similarity in the proportions of zero values across both genes and cells (Fig. 2d-e). This held true for both B cells — with 75% of genes containing over 88% zero counts — and RAM cells, which were less sparse, with 75% of genes containing more than 63% zeros. Simulated cellular library size distributions similarly aligned with empirical data, though with fewer low-end outliers for both cell types (Fig. 2F). This discrepancy may stem from the empirical data’s library size distributions diverging from the log-normal model used in our simulations. In such cases, our method allows alternative library size values to be specified to better reflect the data.

**Figure 2:**
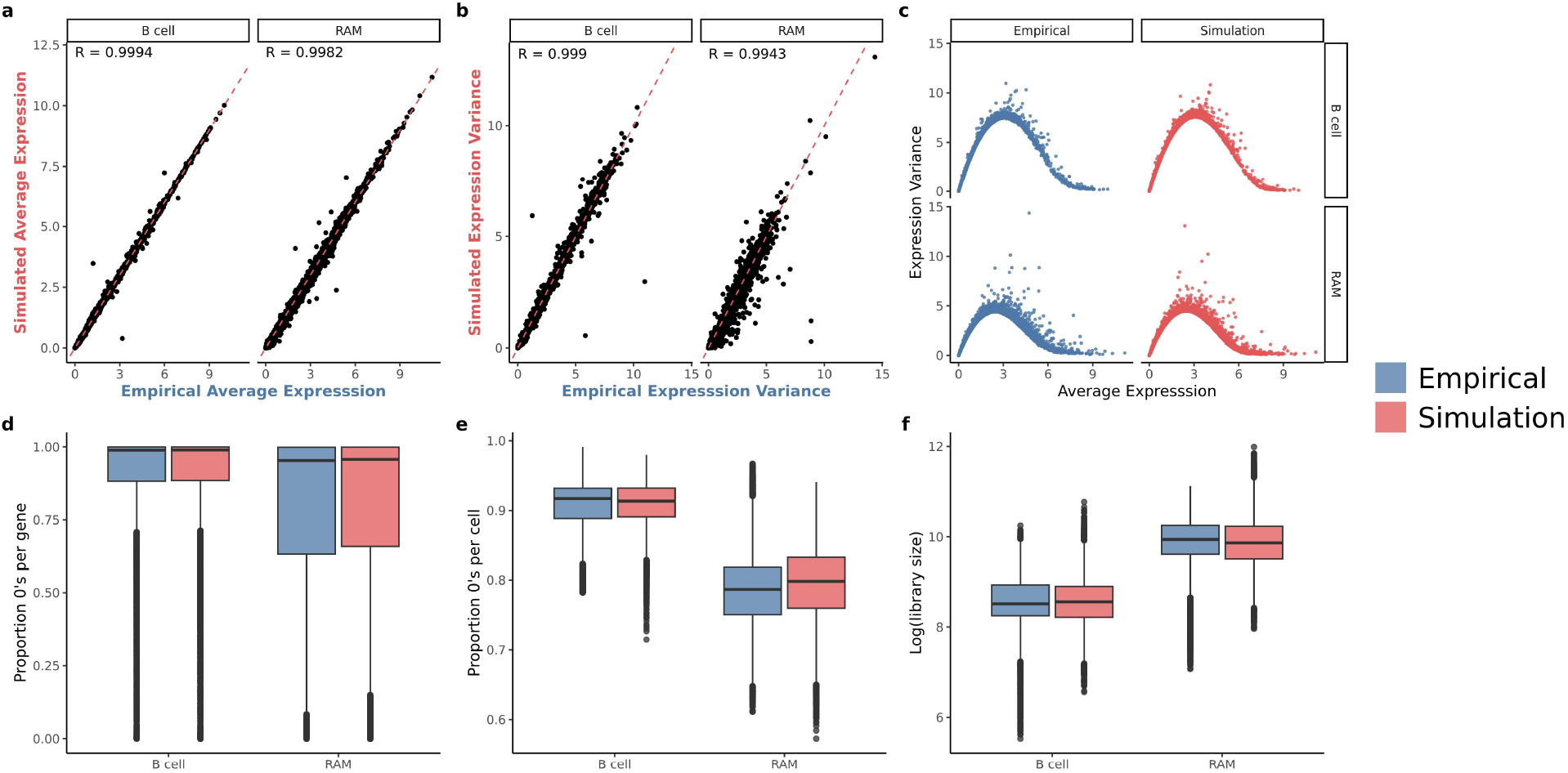
Comparison of metrics for empirical and simulated B cell and RAM data. Metrics include a) average expression, b) variance of expression, c) mean/variance relationship, d-e) proportion of zeros per gene and per cell, and f) cellular library size.

Our simulations also recreated the hierarchical structure of longitudinal scRNA-seq data in RAM cells (Fig. 3) as well as in other cell types as shown in Supplementary material Figs. 3-5). TSNE plots from empirical and simulated data exhibited a similar degree of sample and subject level clustering between cells (Fig. 3A-B). Both subject- and sample-level ICC values exhibited similar distributions for simulated and empirical data (Fig. 3C-D). Additionally, our method replicated the variation in library size distribution across samples in empirical data (Fig. 3E).

**Figure 3:**
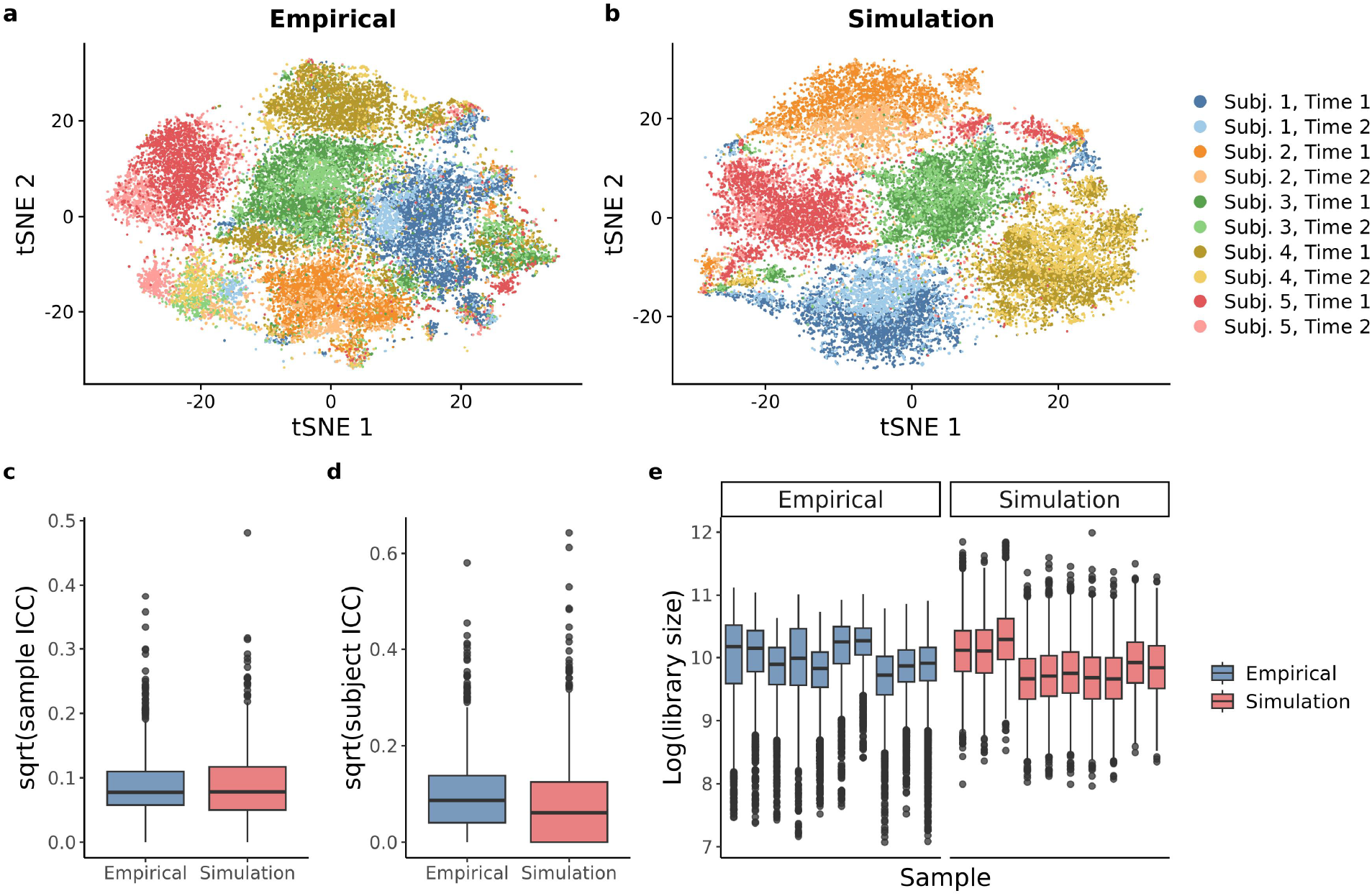
Comparison of sample/subject level variability between empirical and simulated data for RAM data. a-b) tSNE plots from unintegrated data with color representing sample. c-e) Distribution of sample/subject level ICC values and sample-level library sizes.

### 4.2 Power Analysis

Using RESCUE, we were able to perform a power analysis to compare gains in power from recruiting more subjects, gathering data from more timepoints per subject, or sequencing more cells. Increasing the number of subjects, timepoints, or cells in simulated data all led to increased power (Fig. 4). At a 0.05 significance threshold, the average power for simulations with five subjects, 2 timepoints per subject, and an average of 200 cells per sample was 0.41. Increasing the number of subjects had the largest effect on power, as simulated data with 10 subjects had an average power of 0.74 at a 0.05 significance threshold. Increasing the number of timepoints from two to four had the smallest effect on power (average power=0.54 for the 0.05 threshold). Increasing the average number of cells per sample from 200 to 400 had a moderate effect with an average power of 0.65 at a 0.05 threshold.

**Figure 4:**
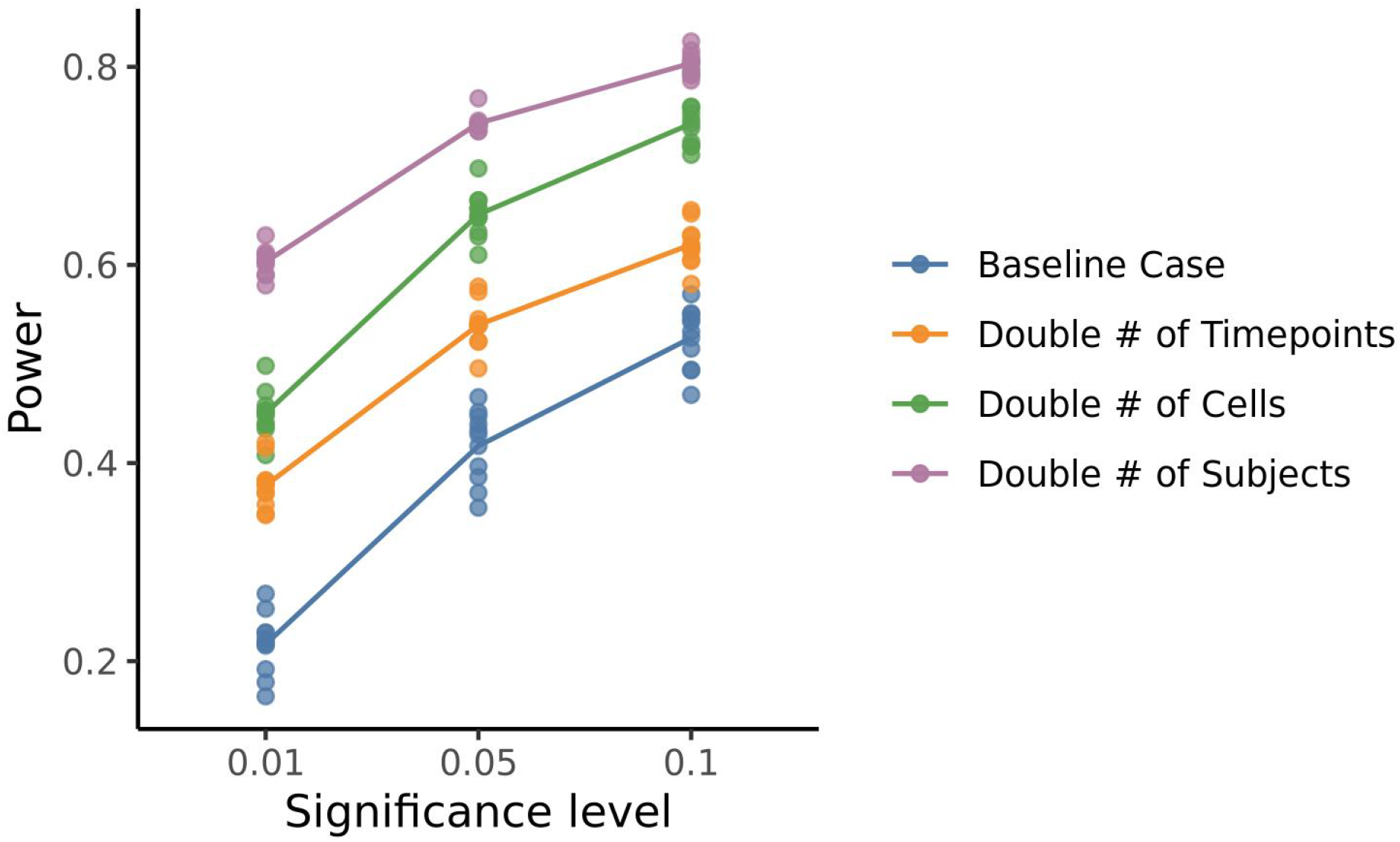
Average power for simulation scenarios at three significance thresholds. Points represent individual simulations.

## 5 Discussion

Single-cell RNA-sequencing is a valuable tool that allows the exploration of gene expression at a cellular level. Longitudinal scRNA-seq experiments enable the study of transcriptional change over time in individual cell types. However, there is not currently guidance on appropriate analysis methods for these types of studies, nor are there resources to assist with study planning. Simulating data that reproduces the complex structure of longitudinal scRNA-seq data is key to effective analysis method evaluation and can also assist in study design.

In this work we established a simulation method, RESCUE, for longitudinal scRNA-seq data and demonstrated that this method could effectively recreate important data properties from empirical data. The mean and variance of normalized gene expression, as well as the distribution of the proportion of zero-count genes across cells and genes, and the cellular library sizes, were similar between empirical and simulated data. We also reproduced the mean-variance relationship exhibited in the empirical data, which is a key characteristic of RNA-seq data. In addition, our simulations captured the hierarchical structure inherent to scRNA-seq data. Simulated data displayed similar clustering patterns between cells from the same subjects or samples in tSNE plots, and the distribution of ICC values was similar at the subject and sample level. Our method also replicated the heterogeneity in library size distribution from sample to sample.

We demonstrated the utility of our method by using simulated data to assess the effect of adding additional subjects, timepoints, or sequencing more cells on the power to detect differential expression across time. In analyzing differential expression in simulated datasets using the `MAST` package with random effects, we found that increasing the number of subjects yielded the greatest increase in power. It should be noted that while we chose to use `MAST` because it allows for control of between-sample and subject variation, this method has not been evaluated or validated for use with longitudinal data. We observed inflated type one error and false discovery rates in our simulations while using this method, and thus additional testing of this method is warranted.

While our simulation framework performed well on six different cell types from two empirical datasets, future research testing the framework on additional cell types would be valuable. Additionally, while our simulation method provides the ability to access and compare analysis methods for differential expression testing in scRNA-seq data, in this work, we did not perform any assessment of these methods. This is an area of research which would be extremely beneficial in advancing the appropriate analysis of longitudinal scRNA-seq studies.

## 6 Availability of Data and Materials

Baseline BAL data from the Mould et al., 2020 dataset is publicly available on the GEO DataSets website under accession number GSE151928. For this study, data from Subjects 2, 4, 5, and 8 were utilized. Data from one other subject, not included in the original GEO dataset, was also used in this manuscript and will be uploaded to the GEO repository under the same accession number prior to publication. Data from Khoo et al., 2023 is available in the GEO database under accession GSE196456.

The RESCUE R package is currently available on github (https://github.com/ewynn610/RESCUE) and we plan to submit it for inclusion in Bioconductor. Code used to simulate and summarize data in this manuscript is available on github at https://github.com/ewynn610/RESCUE_manuscript_code.

