## Supplemental Material for "Simulating Longitudinal Single-cell RNA Sequencing Data with RESCUE"

### 1 Estimating Batch Effect Hyper-Parameters

To estimate the batch variance hyper-parameters $\mu_b$, $\sigma_b$, $\mu_a$, and $\sigma_a$ from empirical data, we first generate a set of genes that is roughly invariant across timepoints or conditions. To identify this set of genes, we aggregate counts for each sample for each gene, and then use the `edgeR` package to fit models for each gene with the condition/timepoint variable as a fixed effect. If the empirical data are longitudinal, subject is also included as a fixed effect. Genes with $\log_2$-fold changes between -0.1 and 0.1 are used to estimate parameters.

Because we calculate $a_{gi}$ and $b_{gij}$ using sample level means, we would expect some level of variation due to sampling error even if there were no true variation in the sample or subject level means. Using central limit theorem we can approximate the degree of variation due to sampling error as:

$$v_{gerror} = \frac{\phi_g * x_g^2 + x_g}{n * x_g^2} \tag{1}$$

where $\phi_g$ is the dispersion for gene $g$, $x_g$ is the global mean expression, and $n$ is the average number of cells per sample/subject. We subtract the gene specific error variance from the total variance to get the variance due to between sample/subject variability which we will refer to as $v^*_{ga}$, $v^*_{ga}$.

Last, to get the parameters, $\mu_b$, $\sigma_b^2$, $\mu_a$, and $\sigma_a^2$, we take the mean and variance across genes for $v^*_{ga}$ and

$v_{ga}^*$. Then, using the relationship between the mean/variance and the log mean and variance parameters of the log-normal distribution, we calculate $\mu_b$, $\sigma_b$, $\mu_a$, and $\sigma_a$. We found that for genes with a high percentage of 0's, the estimates for $v_{ga}^*$ and $v_{ga}^*$ are unstable, so we only use genes with $< 60\%$ 0 counts to estimate the parameters.

### 2 Supplemental Tables and Figures

Table 1: Summary of simulated cell types and batch level hyper-parameters for each cell type

| Cell Type | Dataset | # of Cells | # of Genes | $\mu_a$ | $\sigma_a$ | $\mu_b$ | $\sigma_b$ |
|---|---|---|---|---|---|---|---|
| Cycling | Mould et al., 2020 | 1,540 | 18,196 | -4.68 | 1.14 | -5.27 | 0.99 |
| RecAM | Mould et al., 2020 | 9,041 | 19,410 | -5.06 | 1.21 | -5.58 | 0.91 |
| RAM | Mould et al., 2020 | 21,364 | 26,998 | -4.84 | 1.14 | -5.70 | 0.87 |
| CD4 T cell | Khoo et al., 2023 | 22,362 | 26,871 | -5.75 | 1.73 | -7.01 | 1.50 |
| CD8 T cell | Khoo et al., 2023 | 23,158 | 63,991 | -5.55 | 1.57 | -6.35 | 0.90 |
| B cell | Khoo et al., 2023 | 22,927 | 38,359 | -5.75 | 1.75 | -7.00 | 1.44 |
| NK | Khoo et al., 2023 | 22,362 | 26,871 | -5.50 | 1.54 | -6.35 | 1.00 |

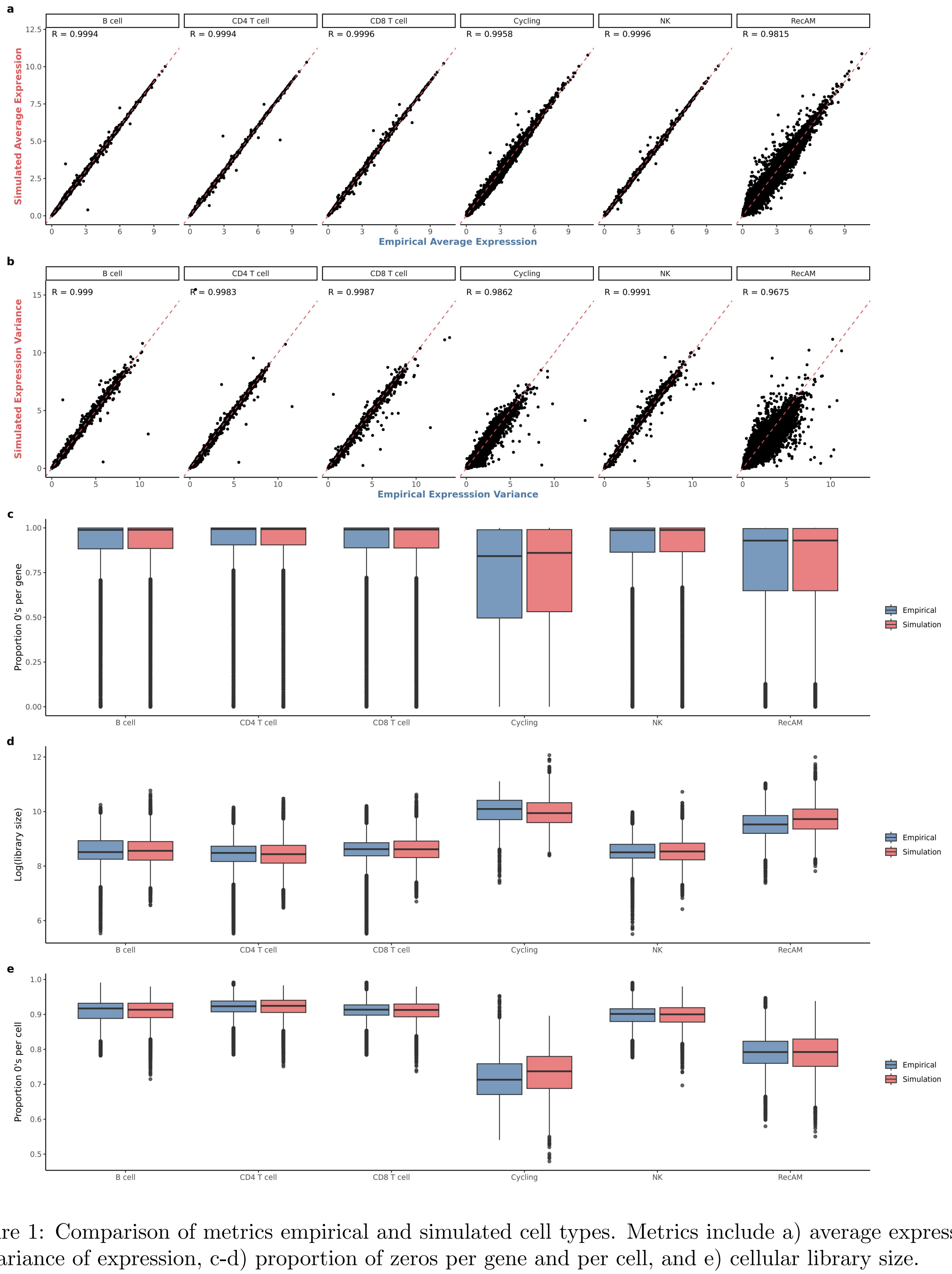


Figure 1: Comparison of metrics empirical and simulated cell types. Metrics include a) average expression, b) variance of expression, c-d) proportion of zeros per gene and per cell, and e) cellular library size.

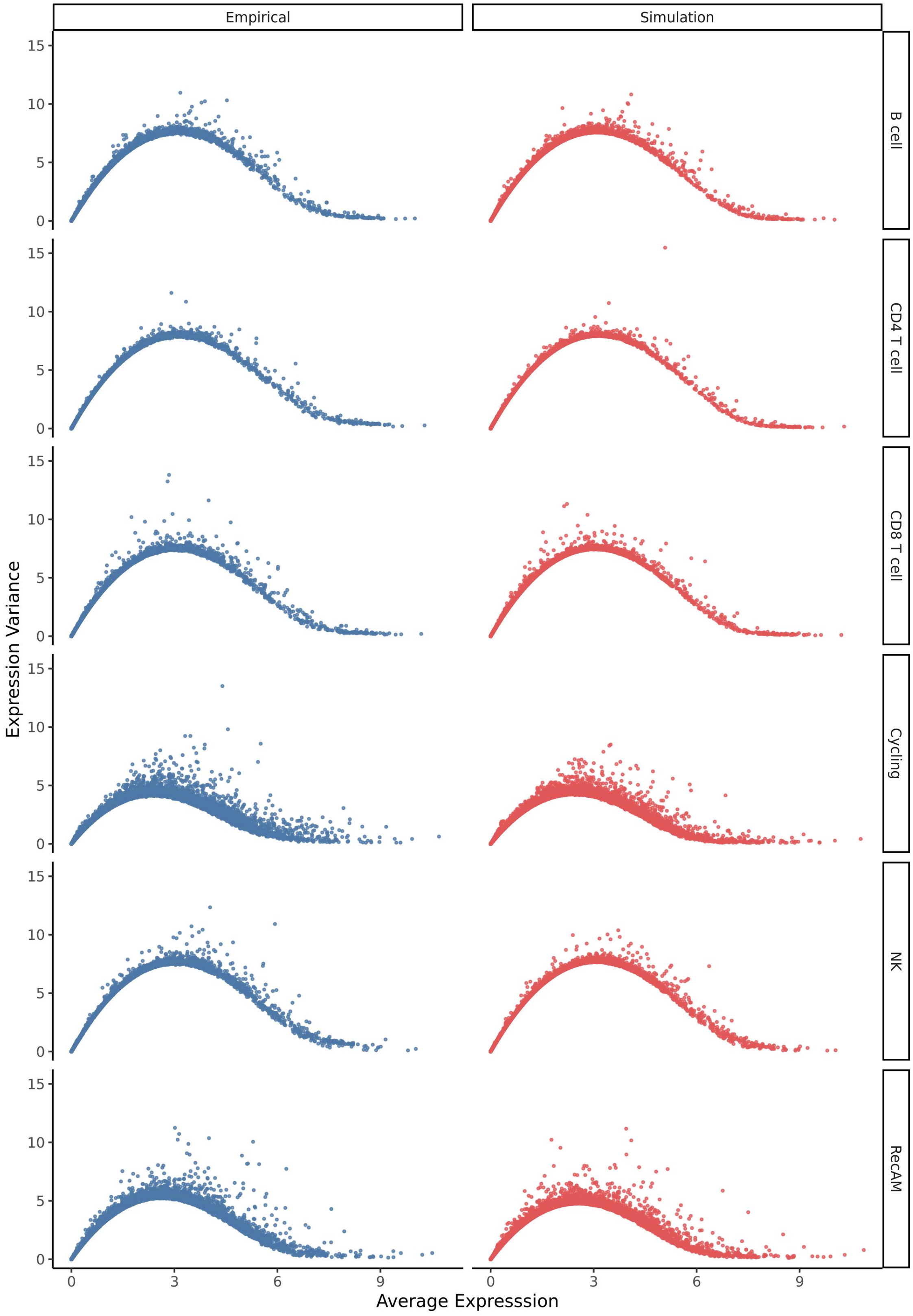


Figure 2: Relationship between mean and variance of gene expression for empirical and simulated data and across cell types.

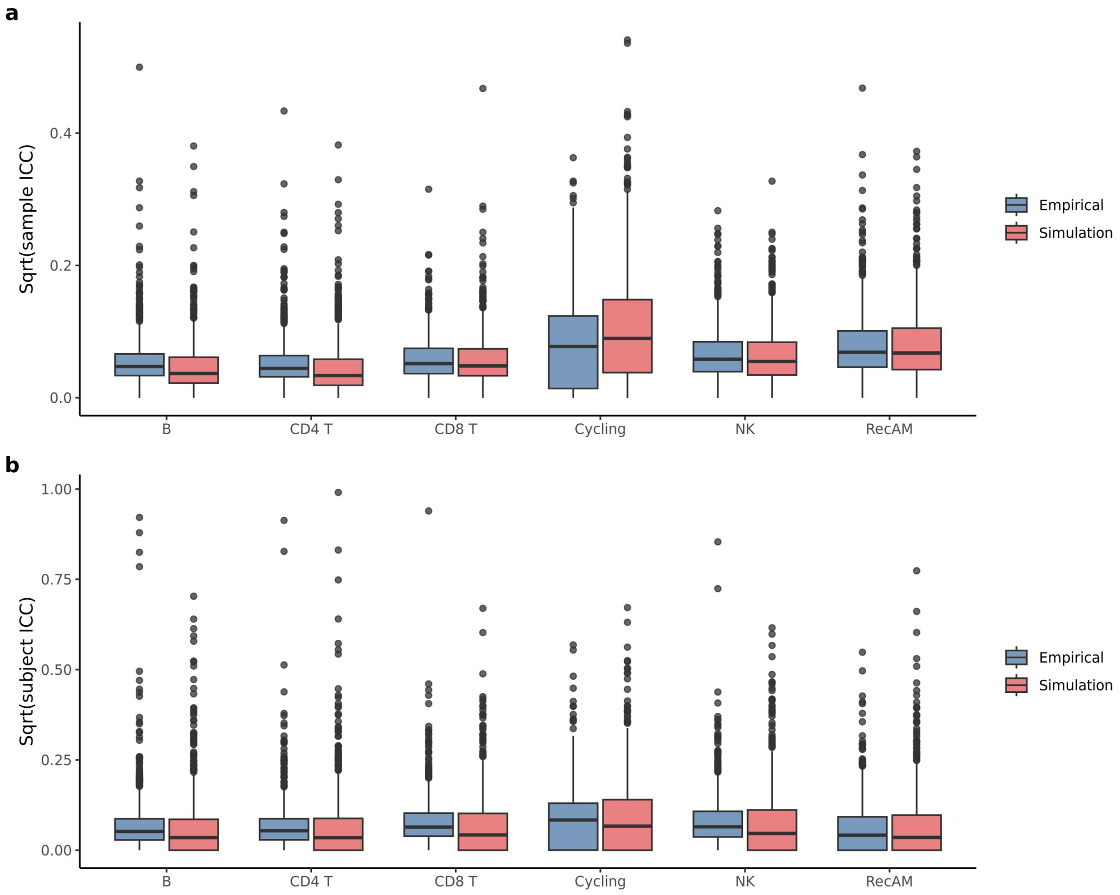


Figure 3: Comparison of distributions of sample level ICC (a) and subject level ICC (b) for all simulated datasets.

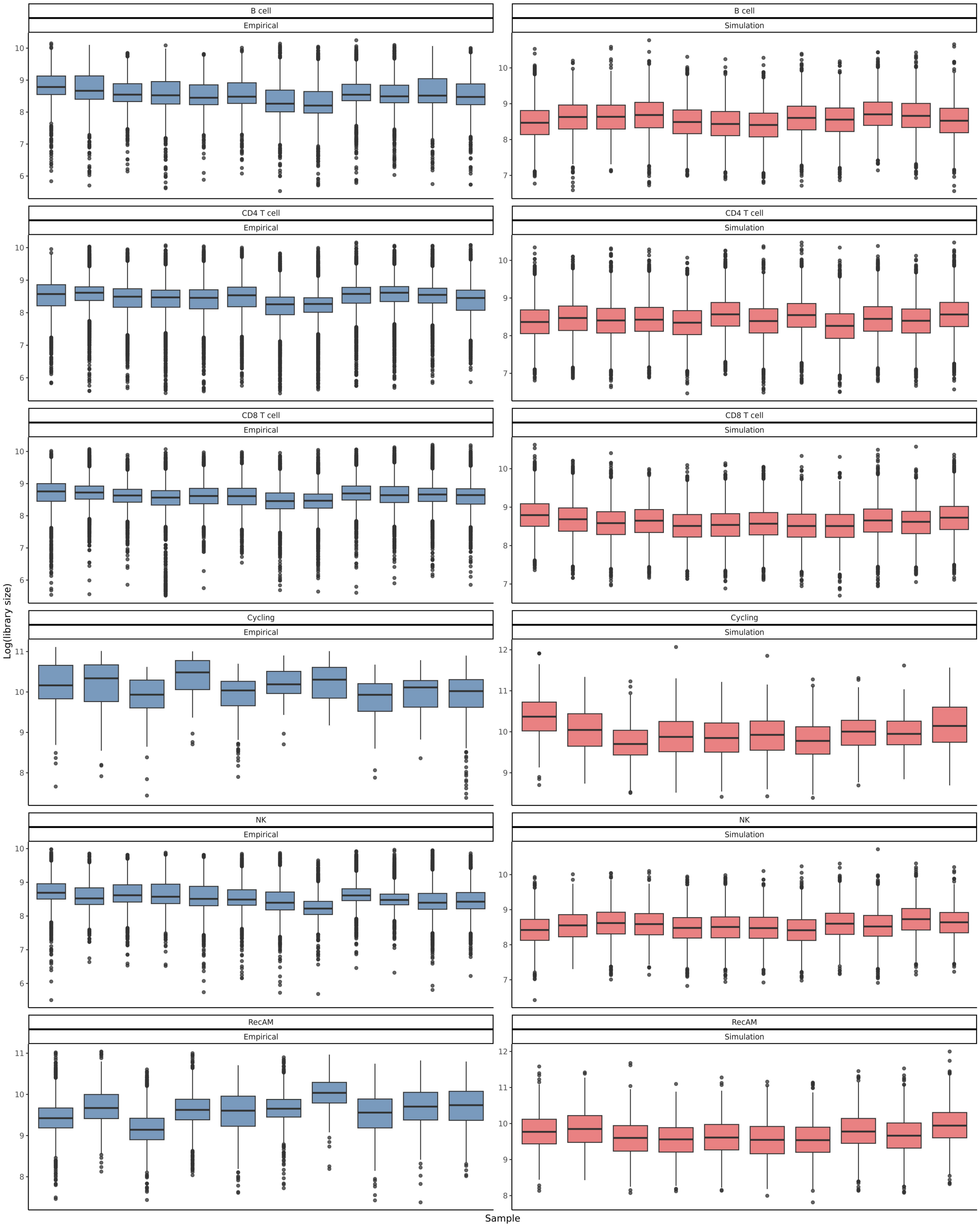


Figure 4: Comparison of sample-level library size distributions for simulated and empirical data and different cell-types.

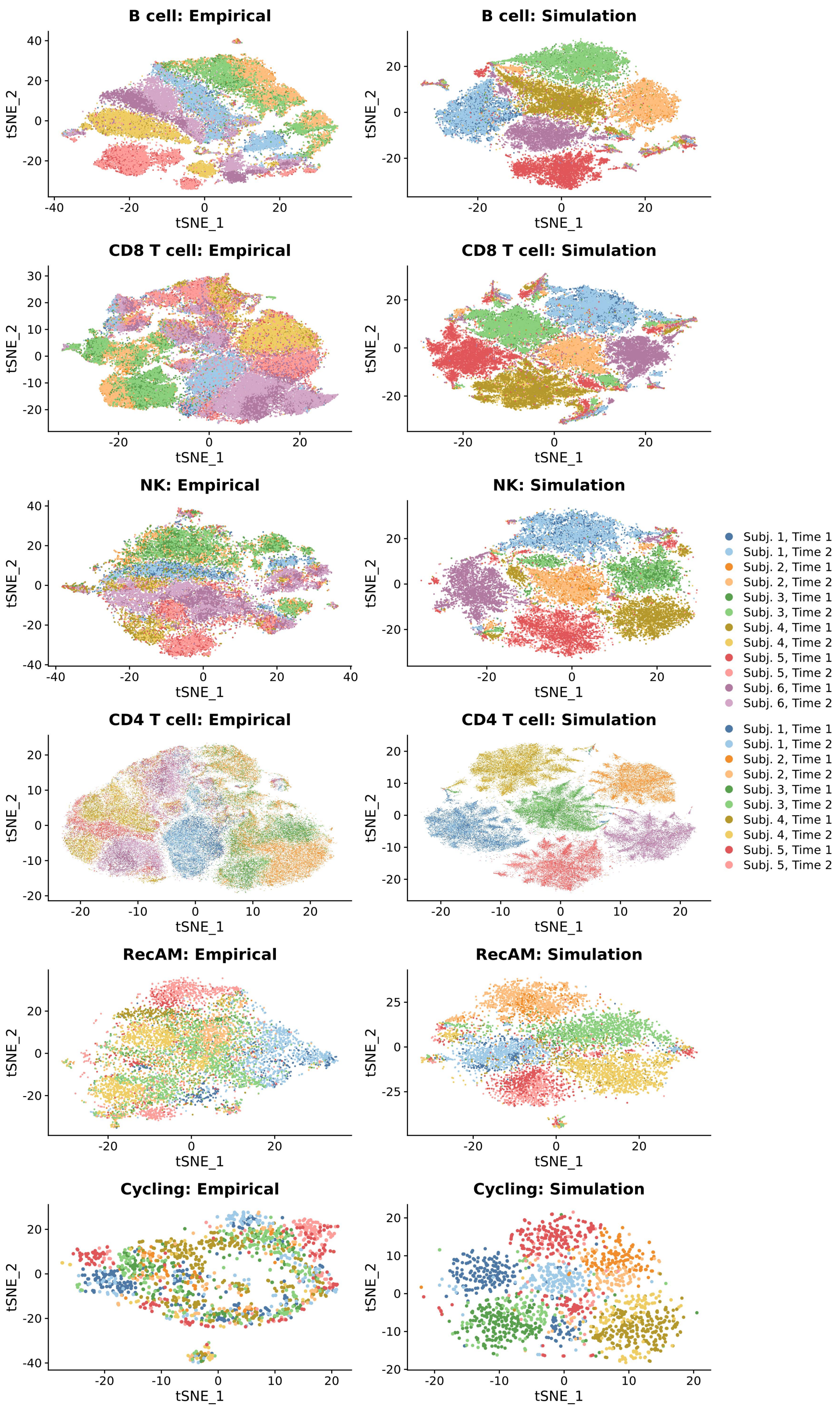


Figure 5: Comparison UMAPS for unintegrated simulated and empirical data and different cell-types. Five random subjects were chosen for the T-cell datasets to avoid overplotting.
